# A device to collect exhaled breath to study biomarkers in small animal models

**DOI:** 10.1101/511931

**Authors:** Yongtao Liu, Yutong Hou, Yuanrui Hua, Youhe Gao

## Abstract

Biomarkers are measurable changes associated with disease. Exhaled breath provides many signs of bodily changes and has been proposed to be a good early biomarker source because it lacks homeostatic mechanisms. Earlier biomarker detection can provide earlier diagnosis, which can bring about more choices and more time for treatment. The benefit of studying exhaled breath in animal models is that most interfering factors can be avoided, and earlier changes in disease may be found. However, there is currently no device that can effectively collect exhaled breath from small animals. In this study, such a device was designed, constructed, and used for the study of candidate biomarkers from the exhale breaths of a papain-induced pulmonary emphysema rat model.

## 1. Introduction

Biomarkers are the measurable changes associated with disease. Exhaled breath accumulates many biological changes and has been proposed to be a good source of early biomarkers because breath lacks the homeostatic mechanisms that maintain bodily stability and remove biomarkers from the blood. The blood, like cerebrospinal fluid, is regulated by the steady-state mechanism of the body, and the components are not easily changed. To maintain normal operation of the brain, the body will transfer away the metabolic waste and substances affecting the brain’s steady state at all costs. The waste is discharged into the blood by direct exchange, and the blood then discharges the waste in the form of urine, sweat, exhaled breath, tears and bile. While urine, exhaled breath, tears, sweat, and saliva are considered bodily waste, they may all be goldmines in the search for early biomarkers. Based on this theory, more biomarker studies should focus on samples discarded from the body, as waste is an essential part of body stability.

For instance, due to non-homeostatic regulatory mechanisms, valuable candidate biomarkers for the very early stages of disease have been found in biospecimen urine [1; 2]. Urine glucose levels were found to be disordered before blood glucose level increases were observed in obese Zucker diabetic rats [3]. In the subcutaneous cancer rat model, the urinary proteome changed significantly before the tumour mass was even palpable. Some of these protein changes have been reported as tumour markers or are associated with tumours [4]. The identification of candidate biomarkers before the occurrence of amyloid plaque deposition has great importance for early intervention in Alzheimer’s disease (AD). Among urinary proteins in 4-, 6-, and 8-month-old transgenic mouse models, 13 are associated with the mechanisms of AD, and 9 have been suggested as AD biomarkers [5]. Pulmonary fibrosis-related proteins are detected in the urine before pulmonary fibrosis formation in rats; if treatment can begin at this time point, then prednisone, which is ineffective in the late stages, can effectively stop fibrosis development [6]. Changes caused by chronic pancreatitis can be reflected in early urinary proteins. New clues for the early diagnosis of chronic pancreatitis have even been found with a small number of animal models [7]. In the urine of bacterial meningitis rat models, differential urinary proteins have also been detected, many of which have been reported as biomarkers of bacterial meningitis in the cerebrospinal fluid and blood [8]. Changes were found in the urinary proteome of rats injected in the brain with C6 glioma cells before tumours were detected by magnetic resonance imaging (MRI). Many of these differential urinary proteins were previously reported to be associated with glioma [9].

As a biomarker source, breath is complementary to urine. The water-soluble waste molecules in the blood are more likely to be removed in the kidney, while gaseous wastes are more likely to be removed in the lung. Disease biomarkers may be discharged as at least one type of waste, or both [10]. The difference between exhaled breath and urine is that the exhaled breath does not linger, like how urine pauses in the bladder. The exhaled breath is generated at any time and is immediately discharged at any time. The advantage of breath over urine is that it can be easily collected at any time, even when oliguria or anuria occur. Most of the gaseous wastes are small molecule compounds (molecular weights less than 200), which can easily pass through the alveolar wall into the exhaled breath and are also known as volatile organic compounds (VOCs). Even after drinking a small amount of alcohol, ethanol can easily be detected from the breath. Therefore, there is reason to believe that exhaled breath may also be able to reflect early changes in disease and have the same potential as urine for earlier and more sensitive detection. Despite the advantage of exhaled breath as a better biomarker source, exhaled breath biomarker research can be impeded by the fact that changes in breath are too complicated to deduce factors associated with any particular pathophysiological condition, especially in human samples [1]. Although human breath is easier to collect for study than animal breath is, clinical exhaled breath samples are difficult to analyse because of the influence of various factors, such as diet and drugs, which made the study of exhaled human breath samples very difficult to explain. The application of animal models is an important means to studying early biomarkers of disease. The advantage of the animal model is that the diet, water and environmental factors are very well controlled. All the differences reflected in breath are directly associated with the disease. The samples can be taken very early since the starting point of the disease is known. Pathological examinations can be used to track disease progression in animal models, which is clinically impossible. The changes in breath found in animal model studies can then be validated in studies with clinical samples.

At the present, there are only a few studies based on animal models. One of the problems is that exhaled breath from animals, especially the exhaled breath of rats and mice, is difficult to collect. Albrecht [11] studied the VOCs in the exhaled breath of rats, who were anaesthetized and ventilated with synthetic air via tracheotomy for 24 h. However, anaesthetics could affect the exhaled breath components, just like how the anaesthetics pentobarbital sodium and chloral hydrate have the same effect on the urine proteome [12].

Many candidate biomarkers from breath have been reported. Nitric oxide (NO) is a biomarker that has been reported to be associated with many diseases in exhaled breath, such as asthma [13]. In addition to NO, exhaled breath contains many types of VOCs. These compounds are usually composed of alkane, an olefin, a lower alcohol, an aldehyde, a ketone, and other aromatic hydrocarbons [14; 15]. Exhaled breath can reflect many lung diseases, including asthma [16], chronic obstructive pulmonary disease (COPD) [17], bronchiectasis [18], cystic fibrosis [19], and interstitial lung disease [20–22]. In total, 1840 VOCs were identified from breath (872), saliva (359), blood (154), milk (256), skin secretions (532), urine (279), and faeces (381) in apparently healthy human individuals [23].

In this paper, we designed a device to collect the exhaled breath of small animals under natural conditions, to minimize the VOCs released from the skin while eliminating the effects from faeces and urine. Subsequently, the device is tested using a rat model of papain-induced emphysema.

## 2. Materials and methods

### 2.1 Exhaled breath collection by a device in an animal model

A device was designed to collect the exhaled breath of experimental animals, without anaesthesia or tracheotomy and in relatively comfortable conditions. The basic goals of this device design were to not only deliver clean air to the animals but also effectively collect the animals’ breaths. The device included air source pipes connected in sequence, and an animal storage box and gas sampling department. The air source was used to supply pure air; the animal storage box was used to hold the animals for exhaled breath collection; and the gas sampling part was used to collect exhaled samples with a Tenax tube and air pump. The incoming air was synthetic air, made by mixing 99.999% high-purity nitrogen and 99.999% high-purity oxygen in a 4:1 ratio (N_2_=79.1%, O_2_=20.9%) (Beijing Chengweixin Industrial Gas Sales Center). The pipe for carrying the gas was constructed with stainless steel or copper, and all the joints were double clip stainless steel joints, which can guarantee air tightness without producing VOCs. The gas flow metre determined the flow rate of the output gas. The animal storage box was constructed of custom glass, and the front end of the box has a tapered structure; the rear end has a cylindrical structure and the conical top opening was connected to the animal’s nose and mouth and then connected to the Tenax tube. The inlet end of the synthetic air pipe was located near the junction of the conical structure and the cylindrical structure to facilitate the animal’s breathing. The area behind the cylinder was used for animal hind limb fixation and excreta discharge. The whole device can eliminate the influence of animal faeces and urine on exhaled breath and minimize the smell from the animal body. The device has a pressure release end, which can release excessive pressure into the environment when the system pressure is too high, thereby reducing the uncomfortableness of the animal being tested. The animal storage box can be designed according to the size of the animals to be tested and in this research, the project designed animal storage boxes in two sizes to be used for collecting exhaled breath from rats and mice.

### 2.2 Establishment of a pulmonary emphysema rat model

To test the feasibility and practical applicability of the device, the study established a pulmonary emphysema rat model. Ten male Sprague-Dawley (SD) rats (180 ± 20 g) were supplied by Beijing Vital River Laboratory Animal Technology Co., Ltd. The animal licence was SCXK (Beijing) 2016-0011. All animals were maintained with free access to a standard laboratory diet, water, and a 12-h light-dark cycle under controlled indoor temperature (22 ± 2°C) and humidity (65-70%) conditions. Animal procedures were approved by the Animal Ethics Committee, Beijing Normal University, and the study was performed according to guidelines developed by the Institutional Animal Care and Use Committee of Beijing Normal University. Papain (12 U/mg) was purchased from Sigma-Aldrich. SD rats were randomly divided into two groups: the control group (n=4) and the experimental group (n=6). The papain-induced pulmonary emphysema rat model was established as follows [24]. SD rats were anaesthetized by intraperitoneal injection with 2% sodium phenobarbital (40 mg/kg), and papain was instilled through the oral cavity and into the trachea at a rate of 3 mg/100 g of animal weight. In the control group, saline was injected at the same volume.

### 2.3 Histological analysis of pulmonary emphysema rats

Four rats in the experimental group and two rats in the control group were randomly sacrificed at 2 days, 4 days, 7 days and 9 days after injection. Lung tissue samples were quickly fixed in 10% formalin. Then, the samples were embedded in paraffin, sectioned, and evaluated with haematoxylin and eosin (H&E) staining and Masson’s trichrome staining.

### 2.4 Breath sample collection and TD-GC-MS analysis

Breath samples of all rats were collected in the device at room temperature. The animal storage box device was always in a state of clean air inflow, and the gas flow rate was controlled at 60 to 80 mL/min. Animals adapted to the environment for 5-10 min after entering the collecting device to become familiar with the environment and eliminate residual waste in the feeding environment. The Tenax-TA tube (a porous polymer 2,6-diphenyl furan resin, 60/80 mesh, 200 mg, Beijing Municipal Institute of Labour Protection) was activated at 300 °C for 2 h in a high-purity nitrogen atmosphere before collection. At the time of collection, the flow rate of the gas pump was controlled at 50 mL/min, the breaths of each animal were collected for 30 min, and approximately 1.5 L of exhaled breath was adsorbed in the Tenax tube. When exhales are collected, the Tenax-TA tube adsorption tube should be placed on ice to allow for Tenax’s adsorption of low molecular weight VOCs. After collection, the sampling tube was plugged with a Teflon stopper, and the information was recorded and stored at 4°C.

The sample with the Tenax-TA tube was thermally desorbed inside a thermal desorption unit with a high-purity nitrogen atmosphere. First, the tube was heated to 250 °C within 5 mins, and simultaneously, the sample was relocated onto a trap at -30 °C. The cold trap was then heated to 250 °C at 100 °C/s, stabilized for 1 min, and the volatiles were transferred to the chromatographic column for separation. A GC capillary column (DB-VRX, film thickness 1.8 μm, inner diameter 0.32 mm, length 60.0 m, Agilent) was used. The sample was stored for a maximum of one week prior to analysis by thermo-desorption-GC-MS (TD-GC-MS, Shimadzu QP2010S). The inlet temperature was 200 °C, and the column oven started analysis at 35 °C, was kept at 35 °C for 10 min, was heated at a rate of 5 °C/min up to 180 °C, then 10 °C/min up to 230 °C, and kept at 230 °C for 5 min. The split ratio was 10:1, the carrier gas was high-purity helium, the interface temperature was 245 °C, the ion source temperature was 250 °C, the acquisition mode was set to scan, and the scanning range was 45 to 200 m/z.

### 2.5 Compound identification and data analysis

Identification of VOCs was based on mass spectrum match (NIST 2008 mass spectral library) with additional confirmation of chromatographic parameters (retention time) to respective pure standards purchased from ANPEL (Shanghai, China). Peak matching and quantification were performed using GCMS Postrun Analysis software (Shimadzu).

## 3. Results

### 3.1. Design of the device

The entire device consists of four parts (**Fig. 1**). The first part is the air source module (**Fig. 1**,10), which provides clean synthetic air for the animal to reduce the impact of pollutants on the sample. The second part is the air transmission and flow rate and pressure control module (**Fig. 1**,20); their main purpose is to further reduce the high-pressure gas released from the gas cylinder (**Fig. 1**,21). The gas flow metre controls the gas flow rate (**Fig. 1**,23), and the stable synthetic air delivered to the animal through stainless steel or pure copper tubing (**Fig. 1**,24). If the pressure in the system is too high, the pressure can be released through the port (**Fig. 1**,22) to stabilize the pressure in the entire system. The third part is the animal module (**Fig. 1**,30). The module body (**Fig. 1**,31) is produced with custom-made glass. It is designed according to the size of the experimental animal (**Fig. 1**,32). The animal head faces inward, and the tail faces outward. In the head of the box, the upper end is the inlet for synthetic air (**Fig. 1**,311), and the front end of the box is the outlet for exhaled breath (**Fig. 1**,312). A small hole (**Fig. 1**,33) is opened on the lower side of the hind limb, and a container is placed underneath for collecting animal faeces and urine. At the leftmost part of the module, there is a small hole (**Fig. 1**,34) where the holder can be vertically inserted to prevent the animal from escaping during the experiment. The fourth part is the exhaled breath collection module (**Fig. 1**,40), wherein the core component is a Tenax-TA tube (**Fig. 1**,41), and the exhaled breath from the experimental animals is enriched as it flows through. The power source of the gas is provided by the constant flow air pump (**Fig. 1**,43). The bubbler can provide a clear view of the gas flow (**Fig. 1**,42). The whole set of equipment does not require electricity (the constant current air pump has a rechargeable battery), which means it is easy to use and has good stability.

**Fig. 1.**
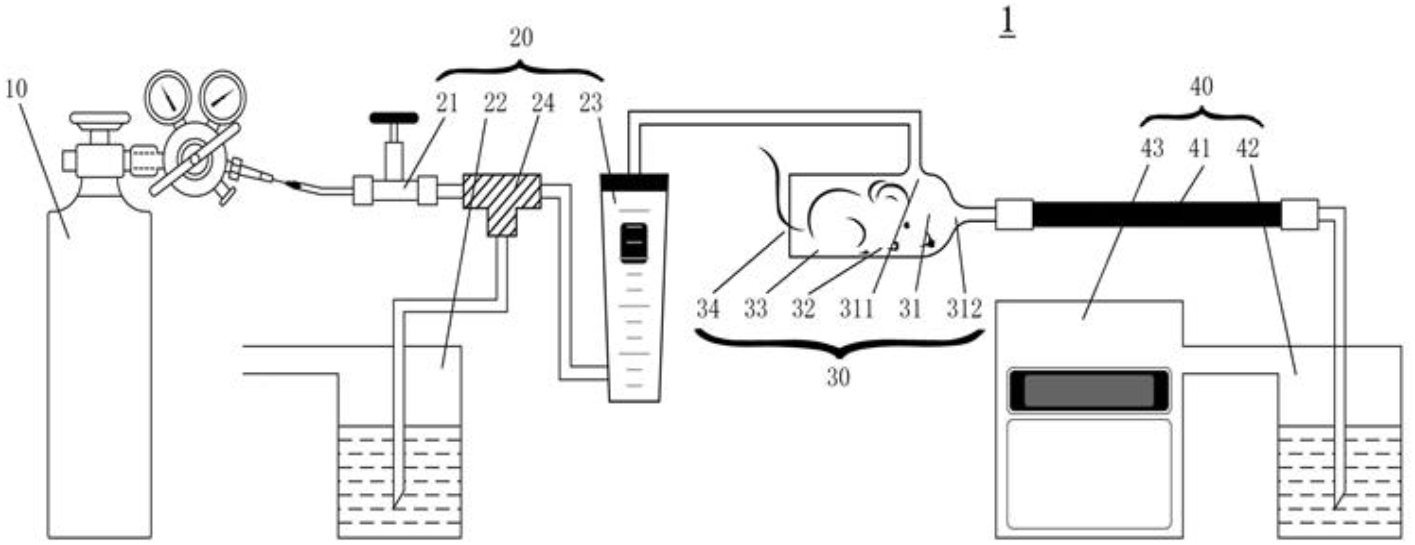
Schematic diagram of the animal breath collection device

The animal storage box is a semi-open environment. When collecting exhaled samples, the animal’s own hair can act as a good barrier to prevent contaminated air outside the device from entering the box and contaminating the collection of exhaled samples (**Fig. 2**).

**Fig. 2.**
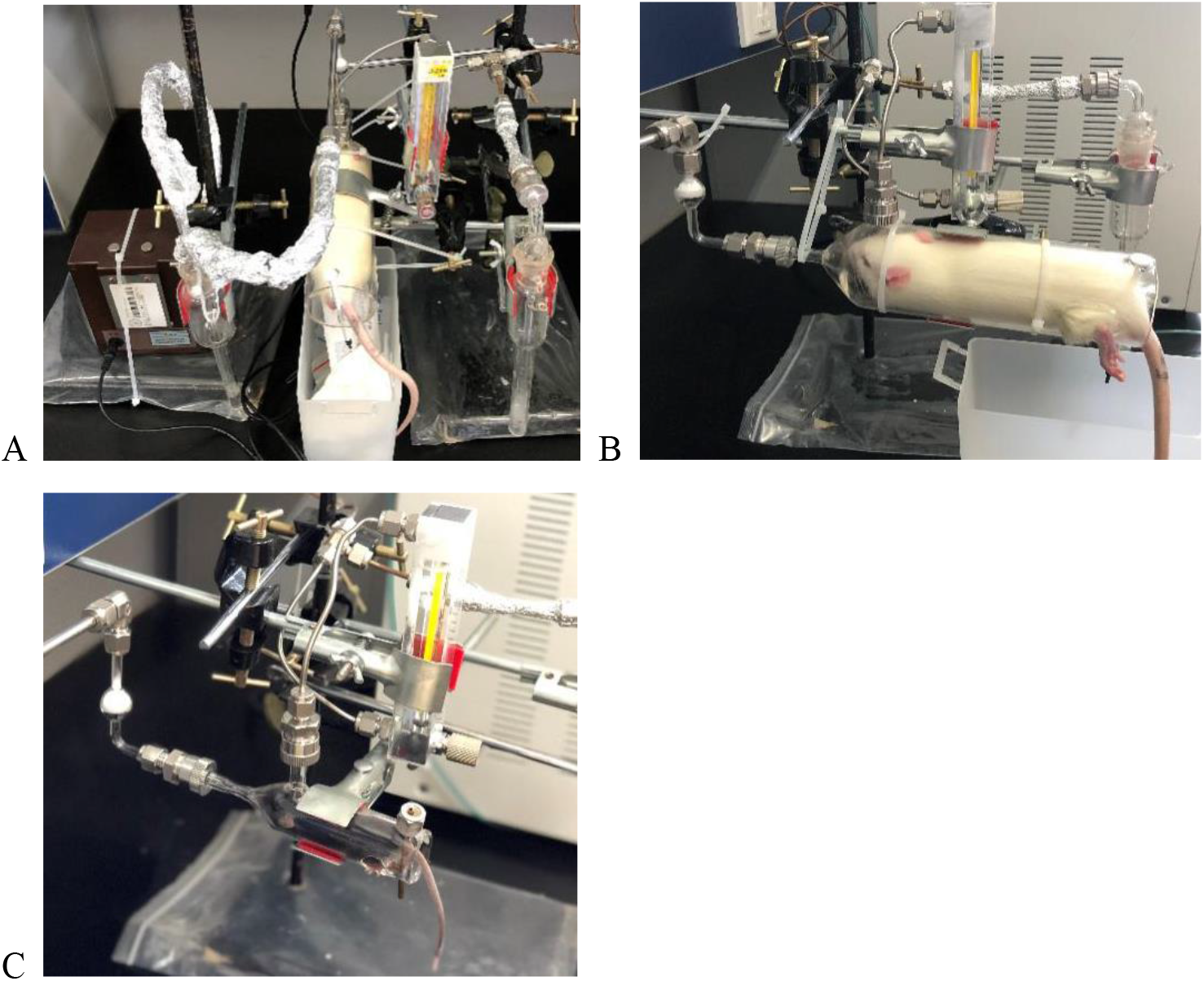
The device is used to collect exhaled breath of experimental animals in a comfortable state. **(A)** SD rat in the device – top view **(B)** SD rat in the device – side view **(C)** C57BL/6 mouse in the device – side view.

Collecting the exhaled breath of small experimental animals is different from that of large animals for which a mask might be used. Small animals can be placed only in certain spaces to collect the air and enrich the exhaled breath. For small animals, the main difficulty in exhaled breath collection and analysis is that exhaled breath is only a small fraction of the container. (**Table 1**). Table 1 shows that the exhaled breath volume of the mice in the unit time was greater than that of the rats. Therefore, when collecting exhaled breath, the exhaled breath collection time of the rats should be appropriately extended.

**Table 1.**
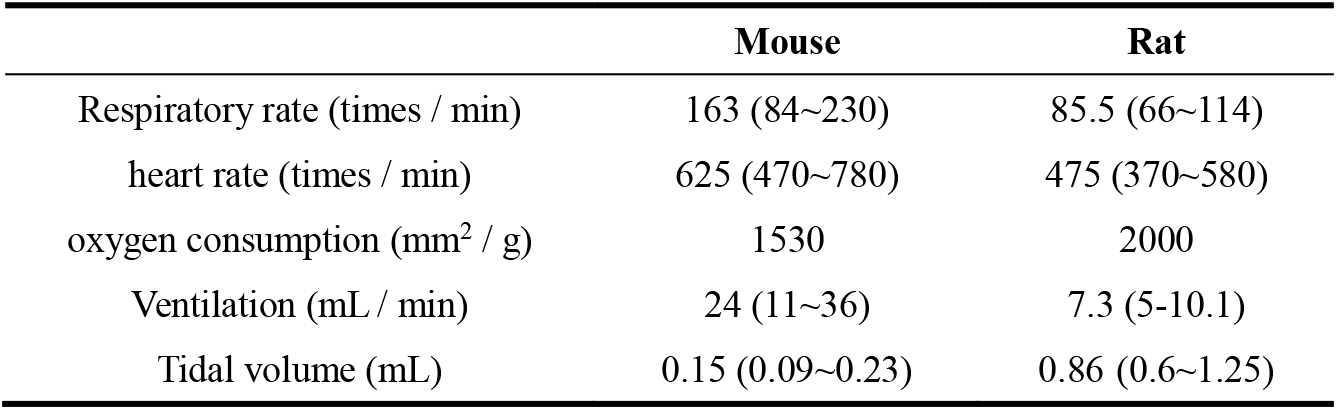
Respiratory-related physiological parameters in rats and mice

|  | Mouse | Rat |
| --- | --- | --- |
| Respiratory rate (times / min) | 163 (84~230) | 85.5 (66~114) |
| heart rate (times / min) | 625 (470~780) | 475 (370~580) |
| oxygen consumption ($\text{mm}^2$ / g) | 1530 | 2000 |
| Ventilation (mL / min) | 24 (11~36) | 7.3 (5-10.1) |
| Tidal volume (mL) | 0.15 (0.09~0.23) | 0.86 (0.6~1.25) |

Notably, part 1 to part 3 of the device can be used separately, and part 4 can be omitted. Replacing part 4 directly connects the device to the online analytical instrument, such as a TD-GC/MS or Q-TOF MS to analyse the VOCs in their exhaled breath. Part 4 is used to store the exhaled breath, and can be suitable for use in some situations where online analysis is not appropriate. Although part 4 can store exhaled breath for a short time, it will likely also lose some component information; at the same time, storage conditions need be considered to prevent contamination of external VOCs [25; 26].

### 3.2 Histopathological changes in the lungs of the pulmonary emphysema rats

H&E staining of the lungs obtained from the pulmonary emphysema rats is shown in **Fig. 3 (A)**. On day 3 after papain administration, the alveolar wall is relatively thin, the alveolar wall is composed of a single layer of epithelium, the size of the alveoli is consistent, and a small amount of connective tissue is seen between adjacent alveoli. As time progressed, the alveolar epithelial stroma gradually thickened and partially broke. On day 14, most of the alveolar walls fused and expanded, the microcapsules of different sizes formed, the alveolar epithelium obviously proliferated, the alveolar space obviously thickened, and a small number of inflammatory cells infiltrated. As shown in **Fig. 3 (B)**, Masson’s trichrome staining revealed very slight pulmonary emphysema in the lungs on day 3, limited lung interstitial emphysema on day 5 and a large number of collagen fibres on day 14. The dramatic changes in the histopathology of the lung tissue suggest that the pulmonary emphysema model was successful and that typical pulmonary emphysema signs appeared 14 days after papain administration.

**Fig. 3.**
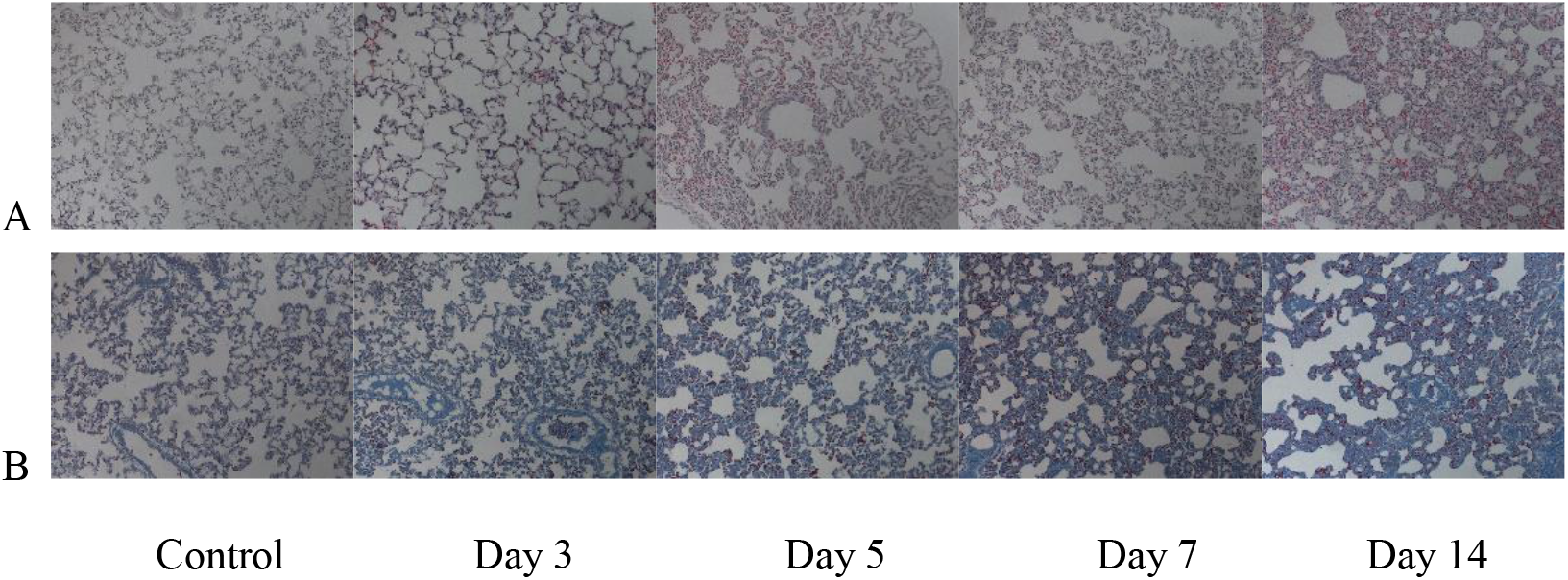
Pathological changes in the lungs of pulmonary emphysema rats. H&E **(A)** and Masson’s trichrome staining **(B)** of rat lungs on days 3, 5, 7 and 14 after papain instillation. The magnification was 200x for the images of H&E and Masson’s trichrome staining.

### 3.3 Analysis of VOCs in the breath of the pulmonary emphysema rats

Through this animal breath collection device, the exhaled breath from rats in this model of emphysema was successfully collected. In the exhaled breath sample, a total of 220 different VOCs were identified (see Supplementary material). The reproducibility of the overall sample analysis is very good. The retention time of each VOC is very stable. When the split ratio is 10:1, the peak width is narrower and the sample analysis effect is better. Among them, 2-ethyl-1-hexanol was a VOC with a significant change, which decreased in the experimental group on day 5 and was still lower than the control group on day 14 when there was an obvious emphysema lesion (**Fig. 4 and Fig. 5**). 2-Ethyl-1-hexanol is an exhaled biomarker that is reported to be closely related to lung cancer [27].

**Fig. 4.**
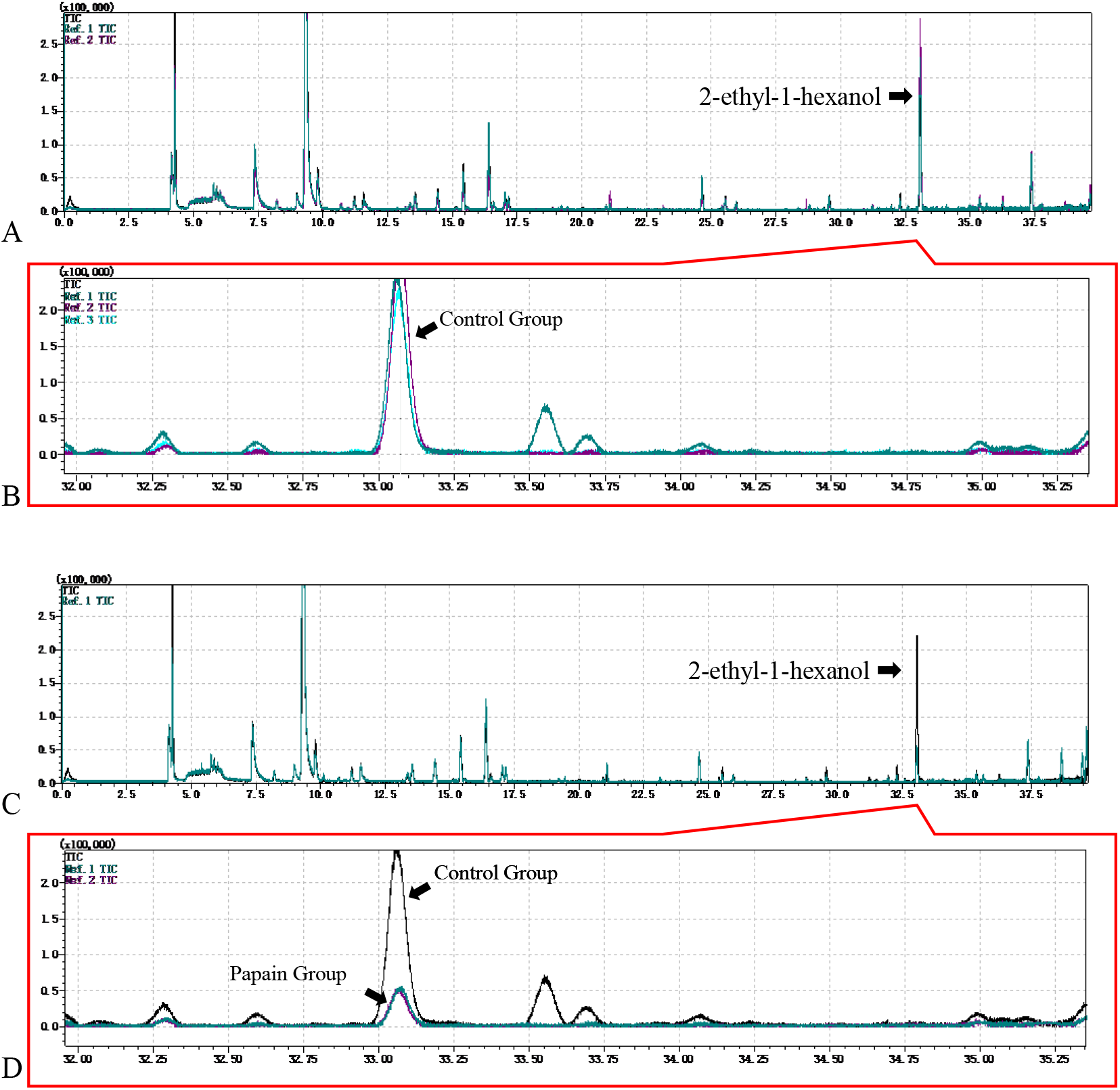
GC-MS spectrum of 2-ethyl-1-hexanol in the exhaled breath of pulmonary emphysema rats. (A) TIC of the control group. **(B)** Enlarged view of 2-ethyl-1-hexanol in the control group. **(C)** TIC of the control group and treatment group. **(D)** Enlarged view of 2-ethyl-1-hexanol in the control group and treatment group.

**Fig. 5.**
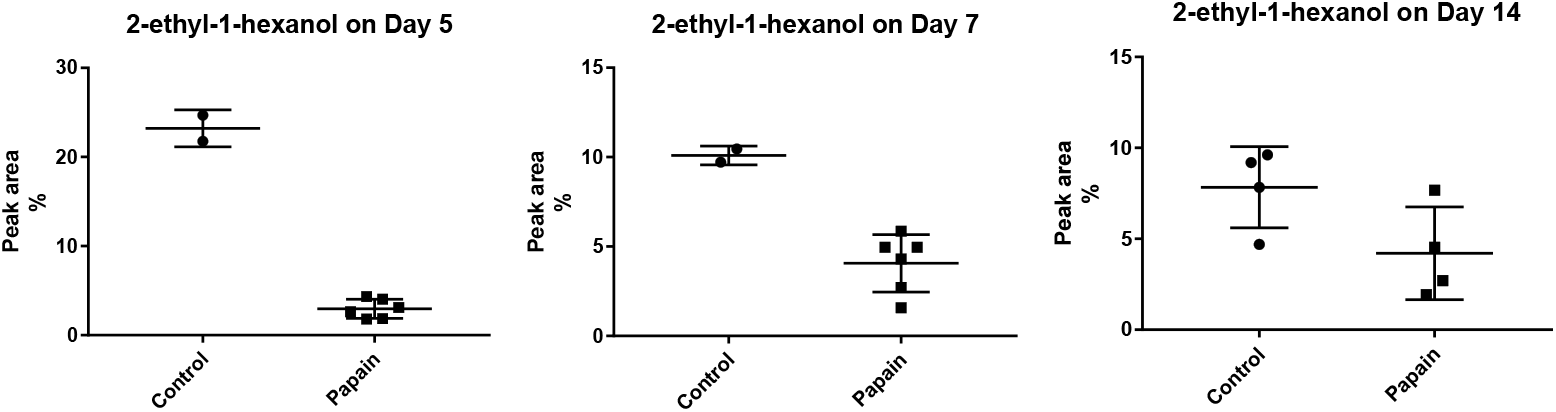
Proportion of the peak area of 2-ethyl-hexanol occupied by exhaled VOCs in a papain-induced pulmonary emphysema rat model at different days (normalized by total peak area).

## 4. Conclusions

This study proposed to discover earlier biomarkers at the small molecular level of exhaled breath in animal models. The use of our breath sample collection system in the animal model was feasible. By establishing a pulmonary emphysema rat model, we showed that it is possible to approximately reveal the difference in exhaled breath during the early stages of the disease.

## Acknowledgements

This work was supported by the National Key Research and Development Program of China (2018YFC0910202, 2016YFC1306300), the Beijing Natural Science Foundation (7172076), the Beijing Cooperative Construction Project (110651103), Beijing Normal University (11100704), and Peking Union Medical College Hospital (2016-2.27).

## Author contributions

Y. Liu and Y. Gao conceived and designed the device. Y. Hou and Y. Hua established the animal model and collected exhaled breath from the model rat. Y. Liu performed the experiments, analysed the data, and wrote the manuscript. All authors approved the final manuscript.

## References

[1] Gao Y 2015 Roadmap to the Urine Biomarker Era MOJ Proteomics Bioinformatics 2

[2] Gao Y 2013 Urine-an untapped goldmine for biomarker discovery? Sci China Life Sci 56 1145–6

[3] Yin W, Qin W W and Gao Y H 2018 Urine glucose levels are disordered before blood glucose level increase was observed in Zucker diabetic fatty rats Science China-Life Sciences 61 844–8

[4] Wu J, Guo Z and Gao Y 2017 Dynamic changes of urine proteome in a Walker 256 tumor-bearing rat model Cancer Med 6 2713–22

[5] Zhang F, Wei J, Li X, Ma C and Gao Y 2018 Early Candidate Urine Biomarkers for Detecting Alzheimer’s Disease Before Amyloid-beta Plaque Deposition in an APP (swe)/PSEN1dE9 Transgenic Mouse Model J Alzheimers Dis 66 613–37

[6] Wu J, Li X, Zhao M, Huang H, Sun W and Gao Y 2017 Early Detection of Urinary Proteome Biomarkers for Effective Early Treatment of Pulmonary Fibrosis in a Rat Model Proteomics Clin Appl 11

[7] Zhang L, Li Y and Gao Y 2018 Early changes in the urine proteome in a diethyldithiocarbamate-induced chronic pancreatitis rat model J Proteomics 186 8–14

[8] Ni Y, Zhang F, An M and Gao Y 2017 Changes of urinary proteins in a bacterial meningitis rat model Sheng Wu Gong Cheng Xue Bao 33 1145–57

[9] Ni Y, Zhang F, An M, Yin W and Gao Y 2018 Early candidate biomarkers found from urine of glioblastoma multiforme rat before changes in MRI Sci China Life Sci 61 982–7

[10] Gao Y 2015 Taking a Deep Breath MOJ Proteomics Bioinformatics 2

[11] Albrecht F W, Huppe T, Fink T, Maurer F, Wolf A, Wolf B, Volk T, Baumbach J I and Kreuer S 2015 Influence of the respirator on volatile organic compounds: an animal study in rats over 24 hours J Breath Res 9 016007

[12] Zhao M, Li X, Li M and Gao Y 2015 Effects of anesthetics pentobarbital sodium and chloral hydrate on urine proteome PeerJ 3 e813

[13] Kharitonov S A, Yates D, Robbins R A, Logan-Sinclair R, Shinebourne E A and Barnes P J 1994 Increased nitric oxide in exhaled air of asthmatic patients Lancet 343 133–5

[14] Fenske J D and Paulson S E 1999 Human breath emissions of VOCs J Air Waste Manag Assoc 49 594–8

[15] Weisel C P 2010 Benzene exposure: an overview of monitoring methods and their findings Chem Biol Interact 184 58–66

[16] Rumchev K, Spickett J, Bulsara M, Phillips M and Stick S 2004 Association of domestic exposure to volatile organic compounds with asthma in young children Thorax 59 746–51

[17] Van Berkel J J, Dallinga J W, Moller G M, Godschalk R W, Moonen E J, Wouters E F and Van Schooten F J 2010 A profile of volatile organic compounds in breath discriminates COPD patients from controls Respir Med 104 557–63

[18] Phillips M, Cataneo R N, Condos R, Ring Erickson G A, Greenberg J, La Bombardi V, Munawar M I and Tietje O 2007 Volatile biomarkers of pulmonary tuberculosis in the breath Tuberculosis (Edinb) 87 44–52

[19] Robroeks C M, van Berkel J J, Dallinga J W, Jobsis Q, Zimmermann L J, Hendriks H J, Wouters M F, van der Grinten C P, van de Kant K D, van Schooten F J and Dompeling E 2010 Metabolomics of volatile organic compounds in cystic fibrosis patients and controls Pediatr Res 68 75–80

[20] Nagaria N C, Cogswell J, Choe J K and Kasimis B 2005 Side effects and good effects from new chemotherapeutic agents. Case 1. Gefitinib-induced interstitial fibrosis J Clin Oncol 23 2423–4

[21] Onozawa M, Hashino S, Sogabe S, Haneda M, Horimoto H, Izumiyama K, Kondo T, Aldana L P, Hamada K and Asaka M 2005 Side effects and good effects from new chemotherapeutic agents. Case 2. Thalidomide-induced interstitial pneumonitis J Clin Oncol 23 2425–6

[22] Chim C S, Ooi G C, Loong F, Au A W and Lie A K 2005 Side effects and good effects from new chemotherapeutic agents. Case 3. Bortezomib in primary refractory plasmacytoma J Clin Oncol 23 2426–8

[23] de Lacy Costello B, Amann A, Al-Kateb H, Flynn C, Filipiak W, Khalid T, Osborne D and Ratcliffe N M 2014 A review of the volatiles from the healthy human body J Breath Res 8 014001

[24] Zhen G, Liu H, Gu N, Zhang H, Xu Y and Zhang Z 2008 Mesenchymal stem cells transplantation protects against rat pulmonary emphysema Front Biosci 13 3415–22

[25] Kang S and Paul Thomas C L 2016 How long may a breath sample be stored for at -80 degrees C? A study of the stability of volatile organic compounds trapped onto a mixed Tenax:Carbograph trap adsorbent bed from exhaled breath J Breath Res 10 026011

[26] Harshman S W, Mani N, Geier B A, Kwak J, Shepard P, Fan M, Sudberry G L, Mayes R S, Ott D K, Martin J A and Grigsby C C 2016 Storage stability of exhaled breath on Tenax TA J Breath Res 10 046008

[27] Jia Z, Zhang H, Ong C N, Patra A, Lu Y, Lim C T and Venkatesan T 2018 Detection of Lung Cancer: Concomitant Volatile Organic Compounds and Metabolomic Profiling of Six Cancer Cell Lines of Different Histological Origins ACS Omega 3 5131–40

